# Fatty acid oxidation drives hydrogen peroxide production by α-ketoglutarate dehydrogenase

**DOI:** 10.1101/2023.11.23.568449

**Authors:** Cathryn Grayson, Ben Faerman, Olivia Koufos, Ryan J. Mailloux

## Abstract

Here, we conducted the first in-depth investigation into sex effects on mitochondrial hydrogen peroxide (mH_2_O_2_) generation in hepatic tissue. Female liver mitochondria produce less mH_2_O_2_ when oxidizing pyruvate, palmitoyl-carnitine, and succinate when compared to male samples. This difference was attributed to superior coupling between fuel metabolism and oxidative phosphorylation (OxPhos) in female liver mitochondria. Examination of mH_2_O_2_ production by individual sites of generation revealed that KGDH was a major source in both male and female liver mitochondria oxidizing pyruvate and malate. Surprisingly, α-keto-β-methyl-n-valeric acid (KMV), a site-specific inhibitor for KGDH, nearly abolished mH_2_O_2_ generation in both male and female liver mitochondria oxidizing palmitoyl-carnitine. KMV did not interfere with the fatty acid oxidation (FAO) pathway and was specific to KGDH. KMV inhibited mH_2_O_2_ production in liver mitochondria from male and female mice oxidizing myristoyl, octanoyl, and butyryl-carnitine. We also supply evidence that KGDH, *not* complex I or complex III, is *the* major mH_2_O_2_ generator in liver mitochondria. Together, we discovered KGDH is a major mH_2_O_2_ source, regardless of sex and during FAO.

## 1. Introduction

The liver is a resilient organ with high regenerative capacity that is integral for whole body energy homeostasis and detoxification reactions (Gonzalez-Teran *et al*, 2016; Wei *et al*, 2018). The Krebs cycle in mitochondria is essential for fulfilling many of these functions as it serves as a point of convergence for metabolic pathways involved in the catabolism and biosynthesis of fatty acids, amino acids, and monosaccharides (Fletcher *et al*, 2019; Satapati *et al*, 2012; Sunny *et al*, 2011). Krebs cycle metabolism is also crucial for supplying reducing power to antioxidant defenses and forming the building blocks for macromolecule biosynthesis (Cappel *et al*, 2019). Pyruvate dehydrogenase (PDH) and α-ketoglutarate dehydrogenase (KGDH) occupy pivotal positions for the entry of carbon into the Krebs cycle as both connect monosaccharide and amino acid metabolism to the electron transport chain (ETC). This connection is formed through the oxidative decarboxylation of pyruvate and α-ketoglutarate, which produce acetyl-CoA and succinyl-CoA, respectively, and NADH (Kiss *et al*, 2013; Soriano-Baguet *et al*, 2023). Acetyl-CoA formed from fatty acid oxidation (FAO) also converges on KGDH since its condensation with oxaloacetate through citrate synthase eventually generates α-ketoglutarate.

It was recently estimated liver mitochondria account for ∼50% of the total H_2_O_2_ in cultured liver cells (Fang *et al*, 2022). However, the reagent used to detect the H_2_O_2_ in this case was Amplex Red, which is not cell permeable, and thus it can be anticipated the amount generated by mitochondria was underestimated. Nevertheless, it has been established mitochondria are major sources of H_2_O_2_. mH_2_O_2_ can be cytotoxic at high enough levels, causing *oxidative stress*, which is defined as an imbalance in oxidants and antioxidants in favor of the former (Sies *et al*, 2017). Lower mH_2_O_2_ doses, however, can have many cell benefits. This led to the conception of two new terms, *oxidative eustress,* and *oxidative distress*, to distinguish between a positive cell interaction with mH_2_O_2_ versus another that is deleterious (Sies & Jones, 2020). The term *eustress* is derived from the Greek word “*eu*” for “*good*” and is achieved when mH_2_O_2_ is maintained at an optimal low concentration of 1-100 nM (Sies *et al*, 2022). Distress is triggered at >100 nM mH_2_O_2_ and results in cell damage and death (Sies *et al*., 2022).

Oxidative eustress pathways are vital for optimal liver health. Liver regeneration is triggered by the mH_2_O_2_-mediated induction of ERK and AKT, stimulating proliferation and survival pathways (Bai *et al*, 2023). mH_2_O_2_-mediated eustress activates NRF2, HIF-1, heat shock factor-1 (HSF-1), promoting antioxidant defense, glycolysis, and protein folding in hepatocytes (Kohli *et al*, 2007; Li *et al*, 2019; Miller *et al*, 2019; Sies, 2017). mH_2_O_2_ also induces cell cycle proteins like cyclin D, promoting cell division in response to a partial hepatectomy (Bai *et al*, 2015). Should an influx in mH_2_O_2_ exceed antioxidant capacity, mH_2_O_2_ will accumulate beyond the eustress range and become cytotoxic (Meng *et al*, 2022; Scialo *et al*, 2020). Specifically in the context of the liver, prolonged distress leads to hepatic pathologies such as metabolic associated fatty liver disease (MAFLD), which begins with non-alcoholic fatty liver disease (NAFLD) and may progress to cirrhosis and hepatocellular carcinoma (Li *et al*, 2023; Ma *et al*, 2023). Although the precise hepatocellular mechanisms for its development are still being delineated, abnormal mitochondrial function causing redox imbalance through the sustained increase in mH_2_O_2_ generation is a common distinguishing feature for the progression of MAFLD (Navarro *et al*, 2017). The distinctive dysfunctions that occur at the mitochondrial level are impairment of fatty acid oxidation and fatty acid overload, abnormal Krebs cycle metabolism, and defects in oxidative phosphorylation (OxPhos), which results in a loss in redox poise and homeostatic control over mH_2_O_2_ (Satapati *et al*, 2016; Talari *et al*, 2023).

Although many benefits of mH_2_O_2_ are established, its sources in mitochondria remain somewhat elusive. Indeed, mitochondria can contain up to 16 mH_2_O_2_ sources. Further, complex I and III of the ETC are often viewed as the main sources of mH_2_O_2_, but this convention has been challenged by the growing evidence showing PDH and KGDH are important mH_2_O_2_ generators. Several studies demonstrated KGDH has a high mH_2_O_2_ production rate in synaptosomes and isolated mitochondria (Horvath *et al*, 2022; Starkov *et al*, 2004; Tretter & Adam-Vizi, 2004). It was reported later that KGDH is 8x more effective at mH_2_O_2_ generation when compared to complex I in muscle (Quinlan *et al*, 2014). PDH was also characterized as an important mH_2_O_2_ source in rat muscle, although it was not as potent as KGDH (Quinlan *et al*., 2014). Purified PDH and KGDH of porcine heart origin or from synaptosomes and KGDH in rodent cardiac tissue can generate large quantities of mH_2_O_2_ during reverse electron transfer from NADH (Ambrus *et al*, 2015; Fisher-Wellman *et al*, 2013; Mailloux *et al*, 2016). In the context of the liver, KGDH accounts for ∼35% of the total mH_2_O_2_ production by liver mitochondria and PDH supplies ∼10%, whereas complex I is a negligible contributor (Oldford *et al*, 2019; Slade *et al*, 2017; Wang *et al*, 2023). Overall, this would suggest KGDH, and to a lesser extent PDH, are potent mH_2_O_2_ generators that may trigger eustress signals.

Mitochondria contain sex dimorphisms in mH_2_O_2_ homeostasis, which is related to fundamental differences in antioxidant defense capacity, overall efficiency of the ETC, proton leak-dependent respiration, and preference for fuel source (Borras *et al*, 2003; Justo *et al*, 2005; Nadal-Casellas *et al*, 2010; Valle *et al*, 2007). However, the impact of sex on the native rates of mH_2_O_2_ generation for individual mitochondrial sources has never been delineated. Here, we employed a mitochondrial substrate and inhibitor toolkit that we have developed to interrogate the individual rates of mH_2_O_2_ generation by components of the ETC, specifically complexes I, II, and III, and KGDH to ascertain if the effect of sex on mitochondrial mH_2_O_2_ homeostasis in the liver is associated with differences in the rates of production by these sites. We report for the first time that the main mH_2_O_2_ source in male and female mouse liver mitochondria is KGDH. In a surprising twist, we also found KGDH is a major mH_2_O_2_ generator during fatty acid oxidation in both sexes. Together, our results show KGDH is an important mH_2_O_2_ source in liver mitochondria, regardless of sex, and that it may serve as *the* oxidative eustress signal induction hub for optimal hepatic health.

## 2. Results

### 2.1 Sex dimorphic effects in mH_2_O_2_ generation are due to better mitochondrial coupling efficiency in female liver mitochondria

The goal of this study was to interrogate sex dimorphic effects in mH_2_O_2_ generation by different sources in hepatic mitochondria fueled by nutrients metabolized through the Krebs cycle, complex II of the ETC, and FAO. **Figure 1** provides an experimental design summary and how sex dimorphisms were interrogated. The toolkit of compounds and inhibitors used to probe rates of mH_2_O_2_ production, enzyme activities, and the states of mitochondrial respiration are also supplied in **Figure 1**. We decided to first interrogate sex differences in liver mH_2_O_2_ generation by examining its rate of production using mitochondria supplemented with pyruvate, succinate, palmitoyl-carnitine, and malate under state 4 respiratory conditions **(Figure 2)**. The rate for mH-_2_O_2_ generation was significantly lower in female liver mitochondria when compared to males (**Figure 2**). We also examined sex effects in the contribution of the individual substrates used to the total amount of mH_2_O_2_ by these samples. The contribution of pyruvate towards the overall rate of mH_2_O_2_ production by hepatic mitochondria collected from the male and female liver mitochondria was similar **(Figure 2**). However, there was a clear sex dimorphic effect in the contribution of succinate and palmitoyl-carnitine to the total amount generated by male and female liver mitochondria. Succinate was a more potent inducer of mH_2_O_2_ production in female liver mitochondria, while fatty acids were more potent inducers in samples collected from male mice. **(Figure 2)**.

**Figure 1:**
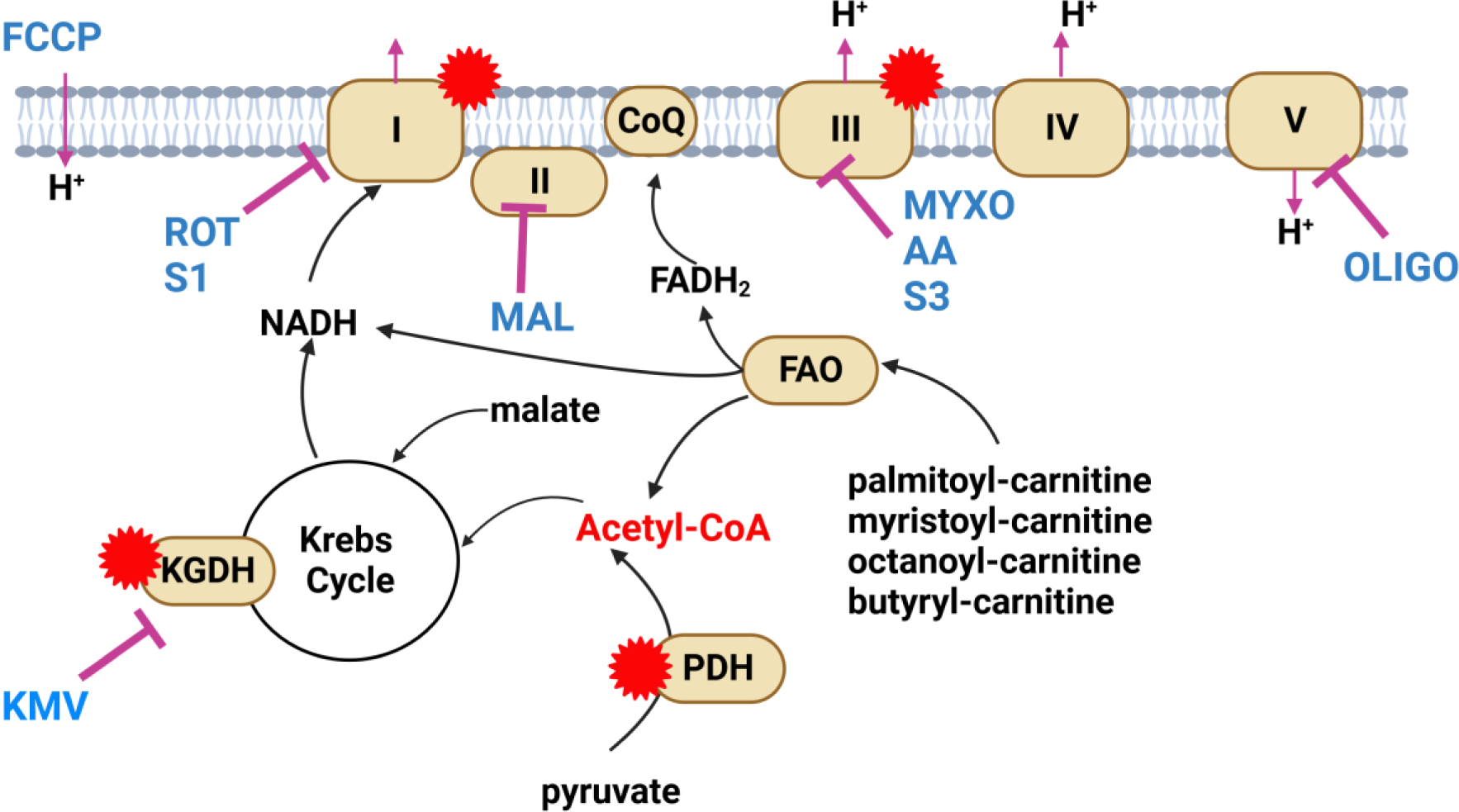
Depiction of the experimental design for interrogating sex dimorphisms in mH_2_O_2_ generation by α-ketoglutarate dehydrogenase (KGDH) and the individual components of the electron transport chain (ETC), complexes I, II, and III. Inhibitor and substrate combinations used for this study are listed in the figure. Reaction mixtures consisted of combinations of pyruvate, succinate, and fatty acyl-carnitines of variable chain length. Malate was added in reactions containing pyruvate or fatty acids to prime the Krebs cycle. Inhibitors used include 3-methyl-2-oxovaleric acid (KMV; KGDH), rotenone and S1QEL 1.2 (Rot, S1; complex I), malonate (Mal; complex II), myxothiazol, antimycin A, and S3QEL 2 (Myxo, AA, S3; complex III), and oligomycin (Oligo; complex V). FCCP was used in the XFe24 assays to estimate the maximal rate of respiration in mitochondria. Different substrates used to power mitochondria for mH_2_O_2_ measurements and O_2_ consumption assays are listed in the figure (pyruvate, malate, succinate, and various fatty acyl-carnitines). The points at which each substrate feeds into nutrient oxidation pathways for mH_2_O_2_ generation and OxPhos is shown. The sources of mH_2_O_2_ and the orientation of production relative to the inner mitochondrial membrane (IMM) is shown as a red star.

**Figure 2:**
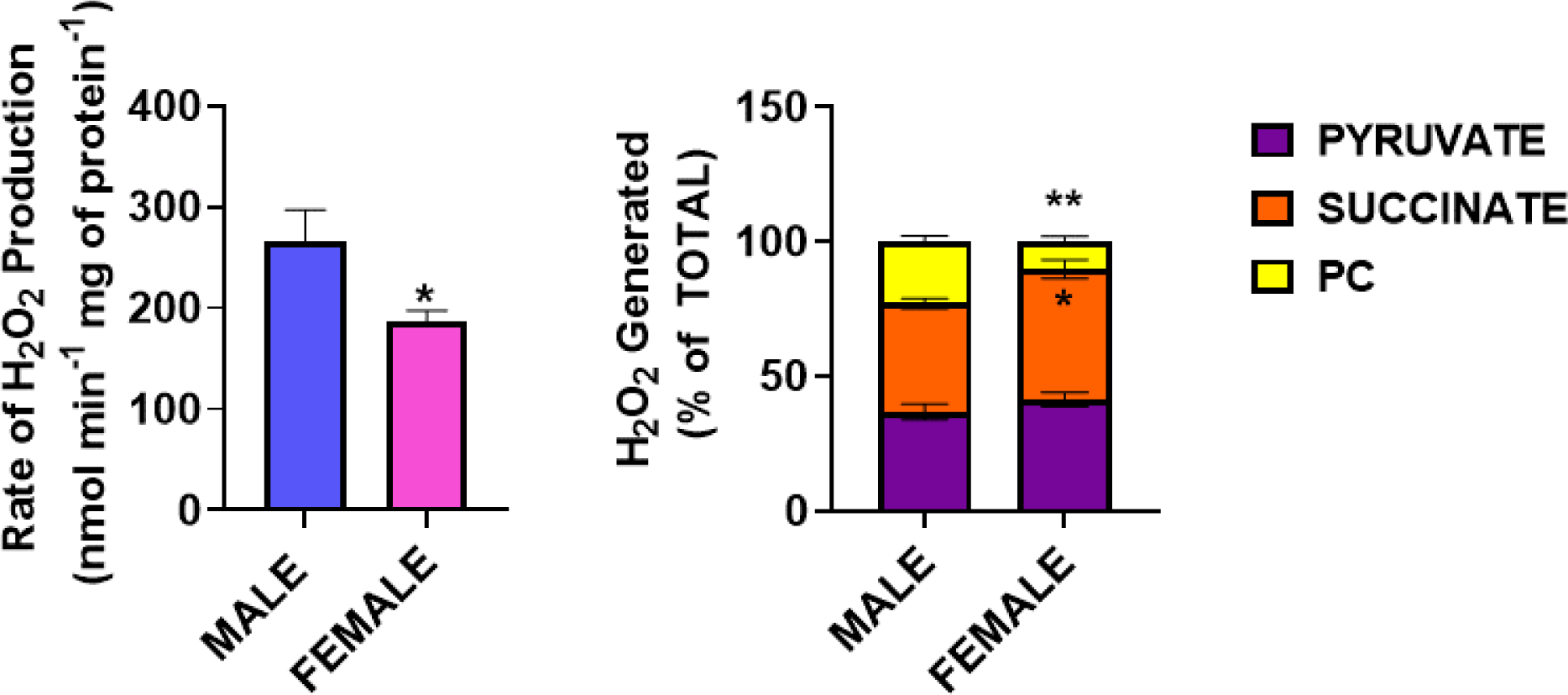
Female liver mitochondria generate less mH_2_O_2_ when compared to male samples. **A.** The rates for mH_2_O_2_ generation by male and female liver mitochondria oxidizing pyruvate, succinate, and palmitoyl-carnitine with malate. N=4, mean ± SEM, two-tailed Student T-Test. **B.** The percent contributions of the different substrates towards the total rate of mH_2_O_2_ production by male and female liver mitochondria. N=4, mean ± SEM, two-tailed Student T-Test.

The lower rate for mH_2_O_2_ generation in female liver mitochondria when compared to males prompted us to examine the bioenergetics of these mitochondria when energized with pyruvate and malate, succinate, or palmitoyl-carnitine and malate. No differences in the O_2_ consumption rate (OCR) under different states of respiration was observed when hepatic mitochondria from male mice were energized with pyruvate and malate, succinate, or palmitoyl-carnitine and malate (**Figure S1**). We made similar findings with the female liver mitochondria **(Figure S1)**. However, the State 3_U_ (activated by injecting FCCP into reaction chambers) in female liver mitochondria was significantly lower when succinate was used as a fuel **(Figure S1)**. Examination of the different states of respiration revealed that liver mitochondria from female mice displayed significantly higher O_2_ consumption rates (OCR) under phosphorylating (state 3) conditions when oxidizing pyruvate and malate, succinate, or palmitoyl-carnitine and malate (**Figure 3A**). Nonphosphorylating O_2_ consumption rates (proton leak dependent or state 4) were also slightly higher in the female liver mitochondria, sometimes being slightly significant in the cases of samples oxidizing pyruvate and malate or palmitoyl-carnitine and malate (**Figure 3A**). No significant differences in the O_2_ consumption rates during state 4_O_ conditions (nonphosphorylating respiration when ADP from state 3 has been exhausted and after the addition of complex V inhibitor, oligomycin) were detected **(Figure 3A**). The exception was the rate for state 4_O_ when pyruvate and malate were the substrates, which was higher in female liver mitochondria **(Figure 3A**). Finally, measurement of uncoupled respiration, or maximal respiration, in the presence of the protonophore, FCCP (State 3_U_), revealed the O_2_ consumption rate under these conditions was ∼2-fold higher in mitochondria collected from female livers **(Figure 3A)**. Calculation of the respiratory control ratio (RCR), a proxy measure for the efficiency of OxPhos revealed it was ∼2-fold higher in females when compared to males in mitochondria oxidizing succinate or palmitoyl-carnitine with malate and ∼2.5-fold greater in female samples energized with succinate **(Figure 3A)**. Next, we measured the overall rate of mH_2_O_2_ generation by mitochondria oxidizing pyruvate and malate, succinate, or palmitoyl-carnitine and malate and plotted these rates against the RCR for the different substrates **(Figure 3B)**. Notably, the rates for mH_2_O_2_ production by female mitochondria oxidizing the individual substrates were found to be lower when compared to the male samples **(Figure 3B)**. There was also an inverse relationship between the RCR value and the rate of mH_2_O_2_ generation. In summary, the higher RCR for the three different substrate(s) accounted for the lower rate of mH_2_O_2_ generation in the female liver mitochondria **(Figure 3B)**.

**Figure 3:**
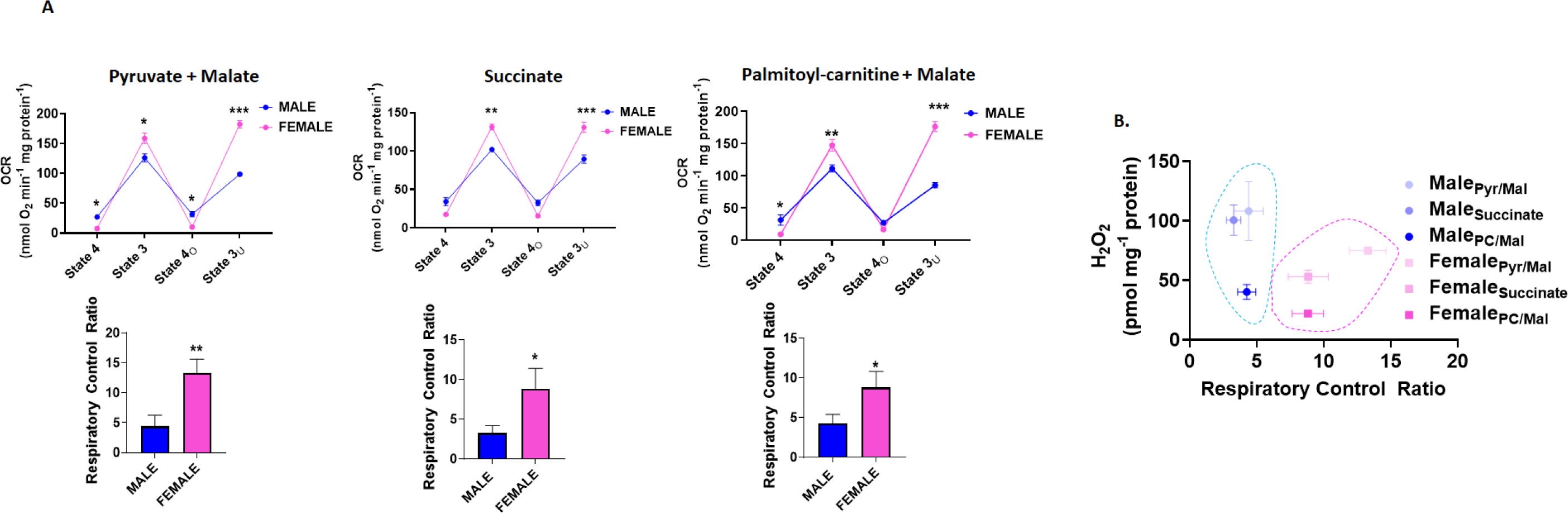
The decreased rate of mH_2_O_2_ production correlates with more efficient nutrient oxidation and OxPhos in female liver mitochondria. **A.** Sequential measurement of rates of O_2_ consumption under state 4, state 3, state 4_O_, and state 3_U_ by male and female liver mitochondria oxidizing different substrates. State 4 was measured first by the addition of just the substrate(s) and then the other states of respiration were measured by the sequential addition of ADP, oligomycin, and FCCP, respectively. All values were corrected for O_2_ consumption recorded after the addition of antimycin A (to arrest the ETC). The values for state 3 and state_O_ were used to calculate the respiratory control ratio (RCR) under each condition N=4, mean ± SEM, two-tailed Student T-Test. **B.** Mitochondria used in the Seahorse XFe24 were used to measure the rate of mH_2_O_2_ generation, which was then plotted as a function of the RCR. N=4, mean ± SEM, two-tailed Student T-Test.

Next, we assessed the expression of KGDH, PDH and several subunits from the ETC complexes to ascertain if the greater coupling efficiency observed in the females was related to protein levels for OxPhos enzymes **(Figure 4)**. The expression of the E1 subunit for KGDH was higher in the female liver mitochondria when compared to males **(Figure 4A)**. The three PDH subunits were also significantly higher when compared to male liver mitochondria **(Figure 4A)**. Use of the OxPhos antibody cocktail revealed no significant differences in SdhB (complex II), UQCRC2 (complex III), MTCO1 (complex IV), or ATP5A (complex V) levels **(Figure 4B)**. However, the abundance of NDUFB8 (complex I) was ∼2-fold greater in female hepatic mitochondria **(Figure 4B)**. Taken together, the more coupling in mitochondrial respiration for ATP production female liver mitochondria account for the sex dimorphism in mH_2_O_2_ production, which is related to the increased expression of PDH, KGDH, and components of the ETC.

**Figure 4:**
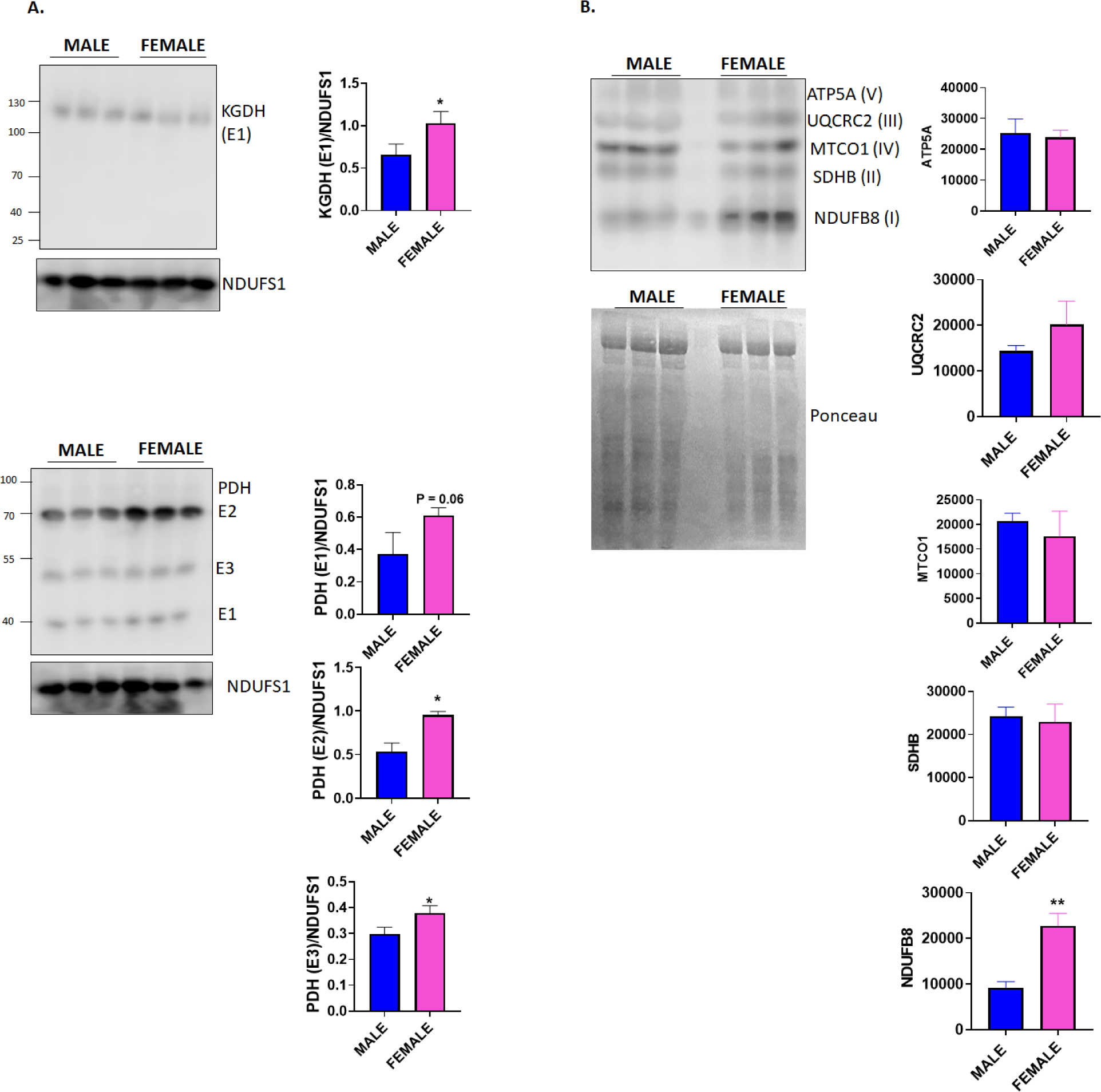
The greater OxPhos efficiency and lower rate for mH_2_O_2_ generation in female liver mitochondria is related to the higher protein expression of PDH, KGDH, and the ETC. **A.** Immunoblot analysis of the E2 subunit of KGDH and the E1-E3 subunits of PDH. NDUFS1 for complex I was used as the loading control. N=3, mean ± SEM, two-tailed Student T-Test. Blots were quantified with ImageJ. **B.** Immunoblot analysis of the expression of the respiratory complexes I-V using the OxPhos antibody cocktail. The subunits that correspond with the individual complexes is indicated in the brackets in the figure. N=3, mean ± SEM, two-tailed Student T-Test. Blots were quantified with ImageJ.

### 2.2 KGDH is a major mH_2_O_2_ source in male and female liver mitochondria

The results above show female liver mitochondria generate less mH_2_O_2_ due to more efficient OxPhos, regardless of the substrate used, and a greater abundance of antioxidant defenses. Next, we examined for sex dimorphic effects on the rates of mH_2_O_2_ generation. We focused our attention on KGDH and the ETC, since we have shown previously these sites account for upwards to ∼90% of the total mH_2_O_2_ formed by male liver mitochondria (Chalker *et al*, 2018; Oldford *et al*., 2019; Slade *et al*., 2017). Titrations with KMV revealed KGDH and downstream generators are major mH_2_O_2_ sources in male liver mitochondria oxidizing pyruvate **(Figure 5A)**. Similar observations were made with hepatic mitochondria isolated from female mice **(Figure 5B)**. Notably, only 0.1 mM KMV was required to induce a robust inhibition of mH_2_O_2_ generation with almost complete abolishment achieved at 0.5 mM in samples collected from males **(Figures 5A and 5B)**. Similar observations were made in females. The rate for mH_2_O_2_ generation by the female liver mitochondria energized with pyruvate was less when compared to the males **(Figure 5C**). Examination of the percent inhibition of mH_2_O_2_ generation by KMV showed the inhibitor almost abolished production in liver mitochondria from both male and female mice **(Figure 5D)**. These findings show KGDH is a major source of mH_2_O_2_ in male and female liver mitochondria oxidizing pyruvate and malate, even though the native rate for mH_2_O_2_ production by is significantly less in female liver mitochondria powered by these substrates.

**Figure 5:**
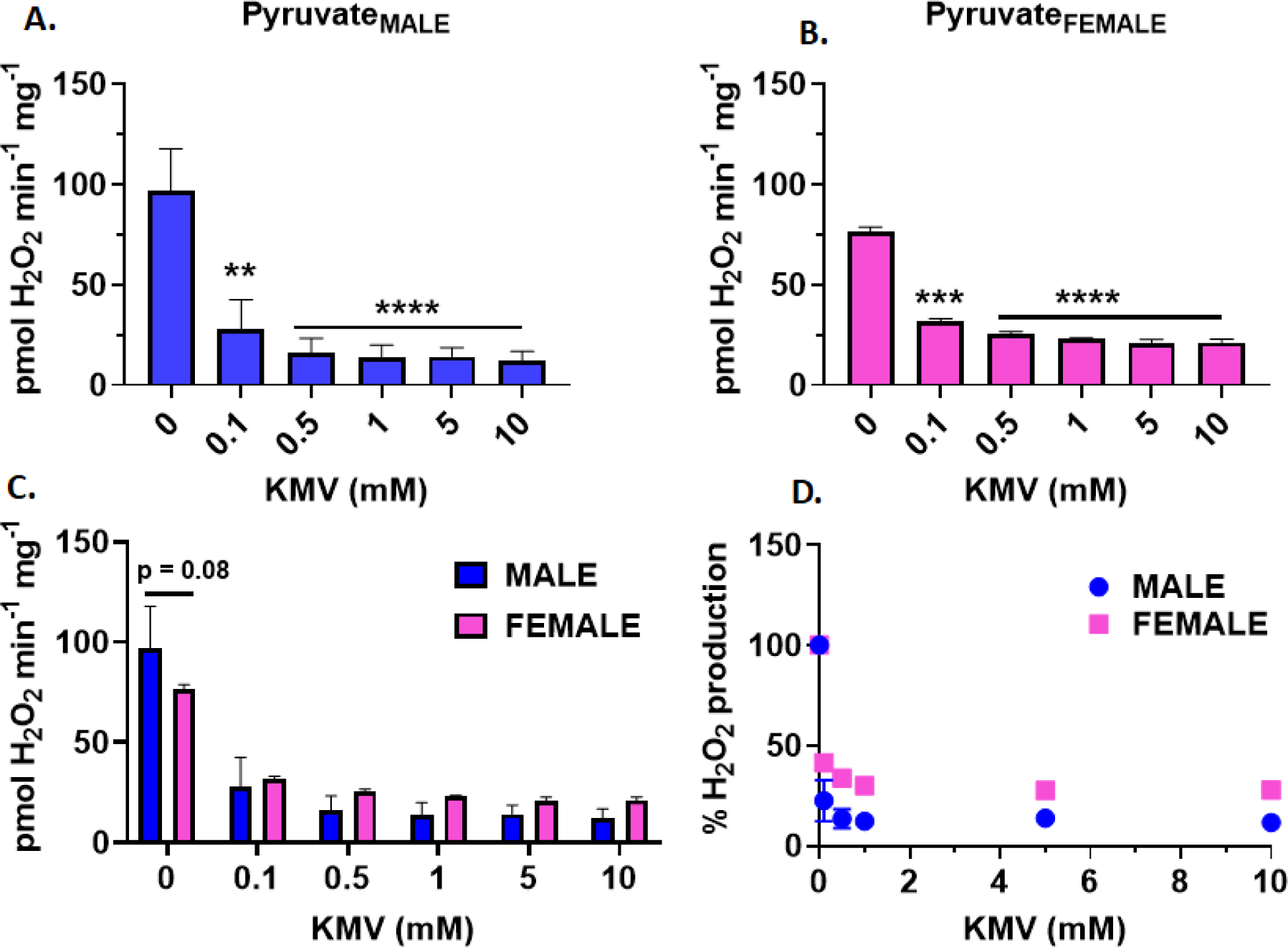
KGDH is a major mH_2_O_2_ supplier in liver mitochondria oxidizing pyruvate and malate. **A.** The effect of increasing doses of competitive and site-specific inhibitor for KGDH, KMV, on the rate of mH_2_O_2_ generation by male liver mitochondria oxidizing pyruvate and malate. N=4, mean ± SEM, 1-way ANOV with a post-hoc Fisher’s LSD test. **B.** The effect of increasing doses of competitive and site-specific inhibitor for KGDH, KMV, on the rate of mH_2_O_2_ generation by female liver mitochondria oxidizing pyruvate and malate. N=4, mean ± SEM, 1-way ANOV with a post-hoc Fisher’s LSD test. **C.** Comparison of the rates of mH_2_O_2_ production by male and female liver mitochondria energized with pyruvate and malate in response and incubated in increasing doses of KMV. N=4, mean ± SEM, 1-way ANOVA with a post-hoc Fisher’s LSD test. **D.** Calculation of the percent change in the rate of mH_2_O_2_ production relative to control in response to increasing doses of malonate. N=4, mean ± SEM, 1-way ANOVA with a post-hoc Fisher’s LSD test.

### 2.3 Mitochondrial succinate oxidation reveals sex dimorphisms in mH_2_O_2_ generation by the ETC

Next, we examined mH_2_O_2_ production during succinate metabolism. For this, all samples were supplied with rotenone to inhibit mH_2_O_2_ generation by complex I through reverse electron transfer (RET) from succinate. **Figure 6A** shows the effect of malonate, a competitive inhibitor for succinate binding to SdhA subunit in complex II, on mH_2_O_2_ by male and female liver mitochondria. Mitochondria were supplied with 5 mM succinate and malonate elicited a strong inhibition of mH_2_O_2_ generation when supplied in equimolar amounts **(Figure 6A)**. The almost complete inhibition of mH_2_O_2_ during succinate oxidation was achieved at 5 mM malonate in female liver mitochondria, whereas double the concentration was required with samples from males **(Figure 6A)**. Comparison of the rates revealed female hepatic mitochondria produced significantly less mH_2_O_2_ when compared to the male counterparts **(Figure 6C)**. However, despite this clear difference in the absolute rates of mH_2_O_2_ production when succinate was oxidized, titration of malonate had a similar inhibitory effect on generation in both male and female liver mitochondria **(Figure 6D)**.

**Figure 6:**
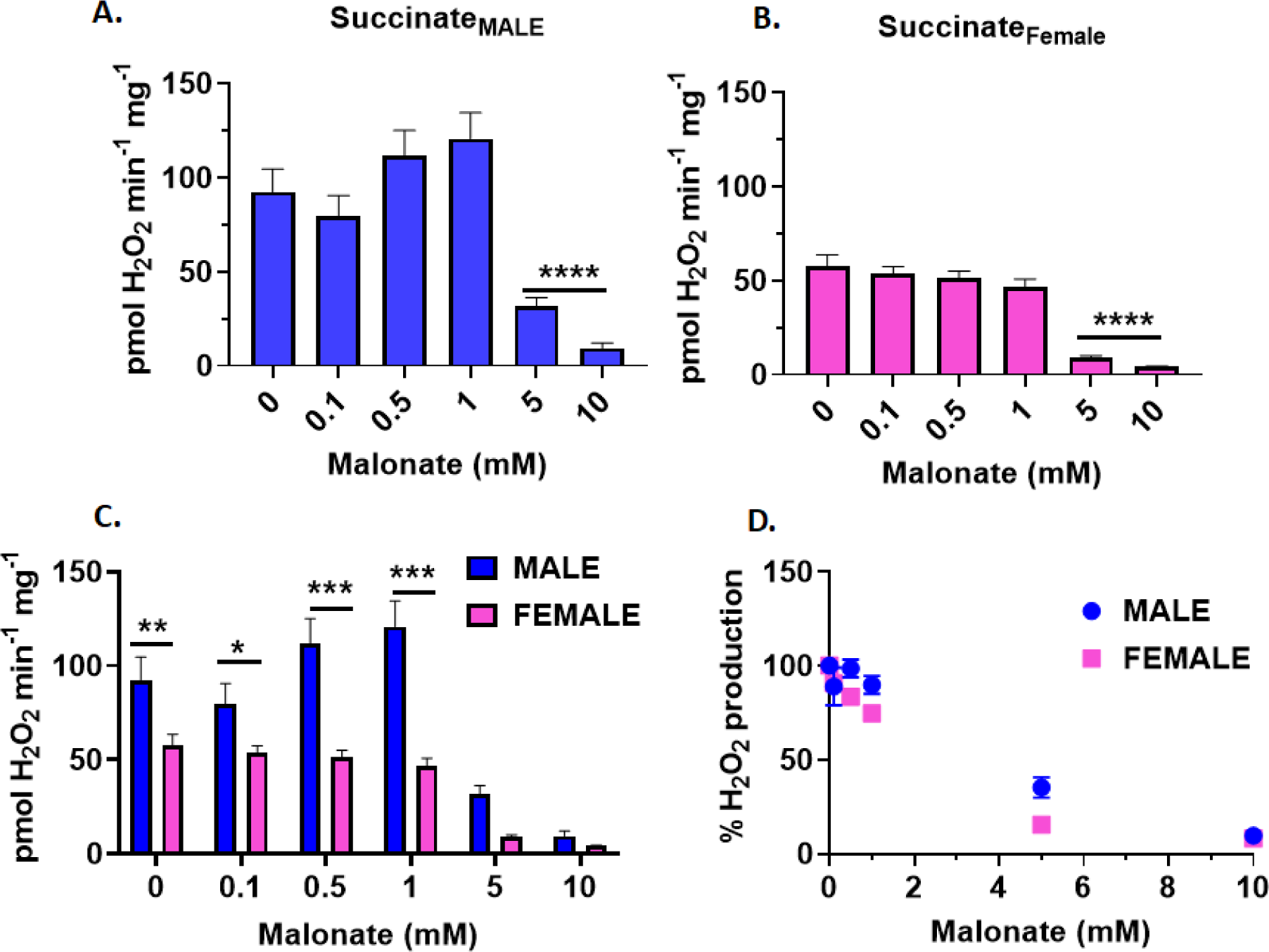
There is no sex dimorphic effect in the response of succinate-driven mH_2_O_2_ production in response to malonate treatment. **A.** The effect of increasing doses of competitive and site-specific inhibitor for complex II, malonate, on the rate of mH_2_O_2_ generation by male liver mitochondria oxidizing succinate. N=4, mean ± SEM, 1-way ANOV with a post-hoc Fisher’s LSD test. **B.** The effect of increasing doses of competitive and site-specific inhibitor for complex II, malonate, on the rate of mH_2_O_2_ generation by male liver mitochondria oxidizing succinate. N=4, mean ± SEM, 1-way ANOV with a post-hoc Fisher’s LSD test.**C.** Comparison of the rates of mH_2_O_2_ production by male and female liver mitochondria energized with succinate in response and incubated in increasing doses of malonate. N=4, mean ± SEM, 1-way ANOVA with a post-hoc Fisher’s LSD test. **D.** Calculation of the percent change in the rate of mH_2_O_2_ production relative to control in response to increasing doses of malonate. N=4, mean ± SEM, 1-way ANOVA with a post-hoc Fisher’s LSD test.

We also examined the impact of myxothiazol on succinate-drive mH_2_O_2_ generation **(Figure 7)**. It Myxothiazol inhibits mO_2_^•-^ by complex III via the limitation of semiquinone radical in its ubiquinol binding site. Thus, myxothiazol is a valuable tool in quantifying the rate of complex III driven mH_2_O_2_ generation. Surprisingly, titrating myxothiazol into reaction chambers did not affect mH_2_O_2_ generation by male liver mitochondria oxidizing succinate **(Figure 7A)**. However, myxothiazol at 5 and 10 µM, respectively, induced a ∼45% decrease in mH_2_O_2_ generation by female liver mitochondria **(Figure 7B)**. Comparing the absolute rates of mH_2_O_2_ production revealed again that female mitochondria generated less when metabolizing succinate, even when myxothiazol was titrated into the reaction chambers **(Figure 7C)**. Analysis of the percentage inhibition of mH_2_O_2_ also confirmed myxothiazol only decreased generation in female mitochondria oxidizing succinate **(Figure 7D)**.

**Figure 7:**
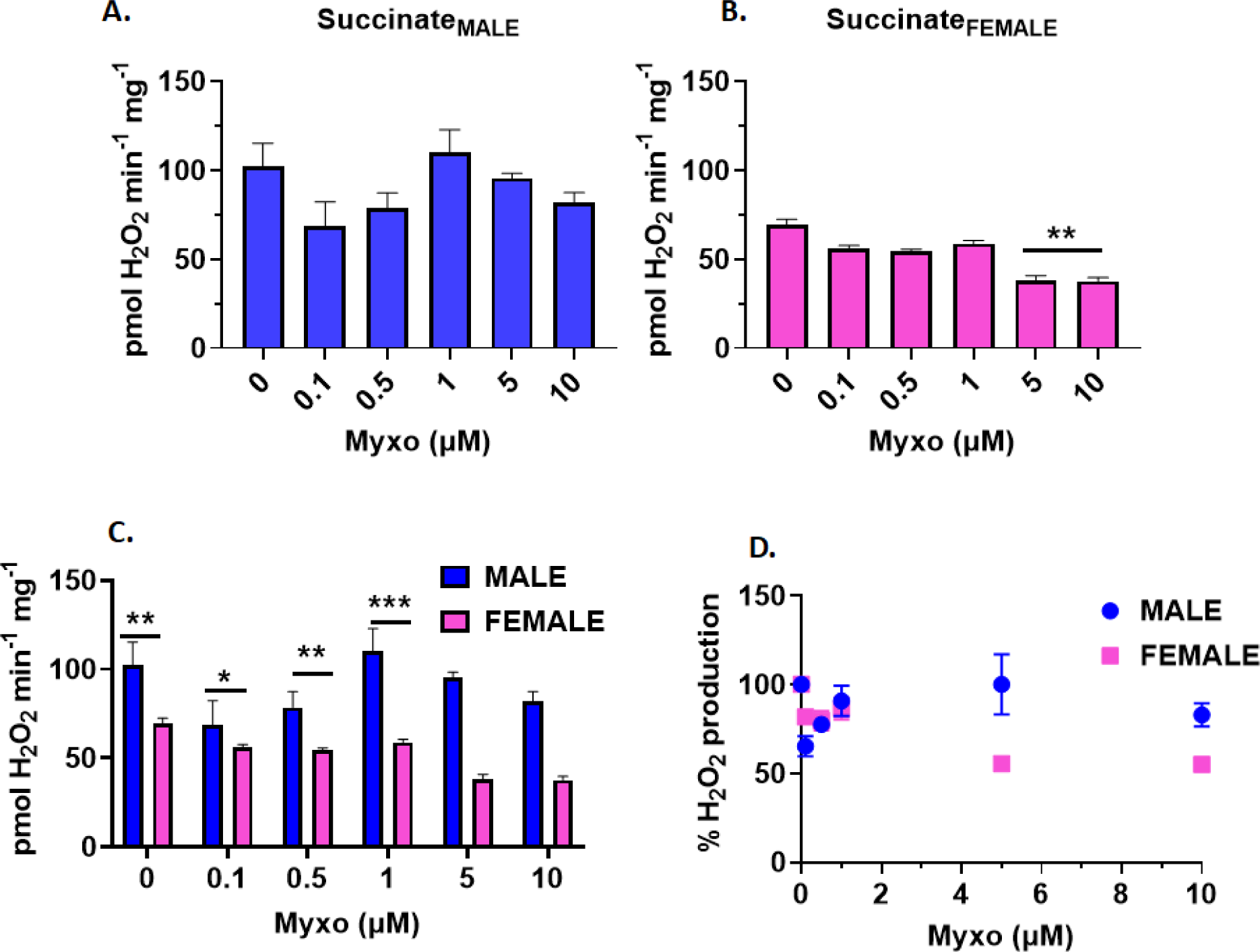
Myxothiazol inhibits succinate-induced mH_2_O_2_ production by female, but not male, liver mitochondria. **A.** The effect of increasing doses of competitive and site-specific inhibitor for complex III, myxothiazol, on the rate of mH_2_O_2_ generation by male liver mitochondria oxidizing succinate. N=4, mean ± SEM, 1-way ANOV with a post-hoc Fisher’s LSD test. **B.** The effect of increasing doses of competitive and site-specific inhibitor for complex III, myxothiazol, on the rate of mH_2_O_2_ generation by male liver mitochondria oxidizing succinate. N=4, mean ± SEM, 1-way ANOV with a post-hoc Fisher’s LSD test. **C.** Comparison of the rates of mH_2_O_2_ production by male and female liver mitochondria energized with succinate and incubated in increasing doses of myxothiazol. N=4, mean ± SEM, 1-way ANOVA with a post-hoc Fisher’s LSD test. **D.** Calculation of the percent change in the rate of mH_2_O_2_ production relative to control in response to increasing doses of myxothiazol. N=4, mean ± SEM, 1-way ANOVA with a post-hoc Fisher’s LSD test.

Next, we examined mH_2_O_2_ generation in mitochondria metabolizing palmitoyl-carnitine. Previous studies established palmitoyl-carnitine generates most of its mH_2_O_2_ through the ETC. Rotenone was added to inhibit mH_2_O_2_ production by RET to complex I. Myxothiazol inhibited mH_2_O_2_ generation by male liver mitochondria oxidizing palmitoyl-carnitine and malate at a concentration of 0.1 µM **(Figure 8A)**. This inhibition remained consistent at higher myxothaizol concentrations. No myxothiazol effect was observed in female liver mitochondria oxidizing palmitoyl-carnitine and malate **(Figure 8B)**. Rates for mH_2_O_2_ generation were significantly higher in male liver mitochondria oxidizing palmitoyl-carnitine and malate when compared to female samples **(Figure 8C)**. Additionally, samples treated with increasing amounts of myxothiazol consistently had the superior rate of mH_2_O_2_ generation **(Figure 8C)**. Examination of the inhibitory effect of myxothiazol as a percentage of the rate of mH_2_O_2_ production in control samples revealed it inhibited mH_2_O_2_ generation in male liver mitochondria by ∼45% at 0.1 µM **(Figure 8D)**.

**Figure 8:**
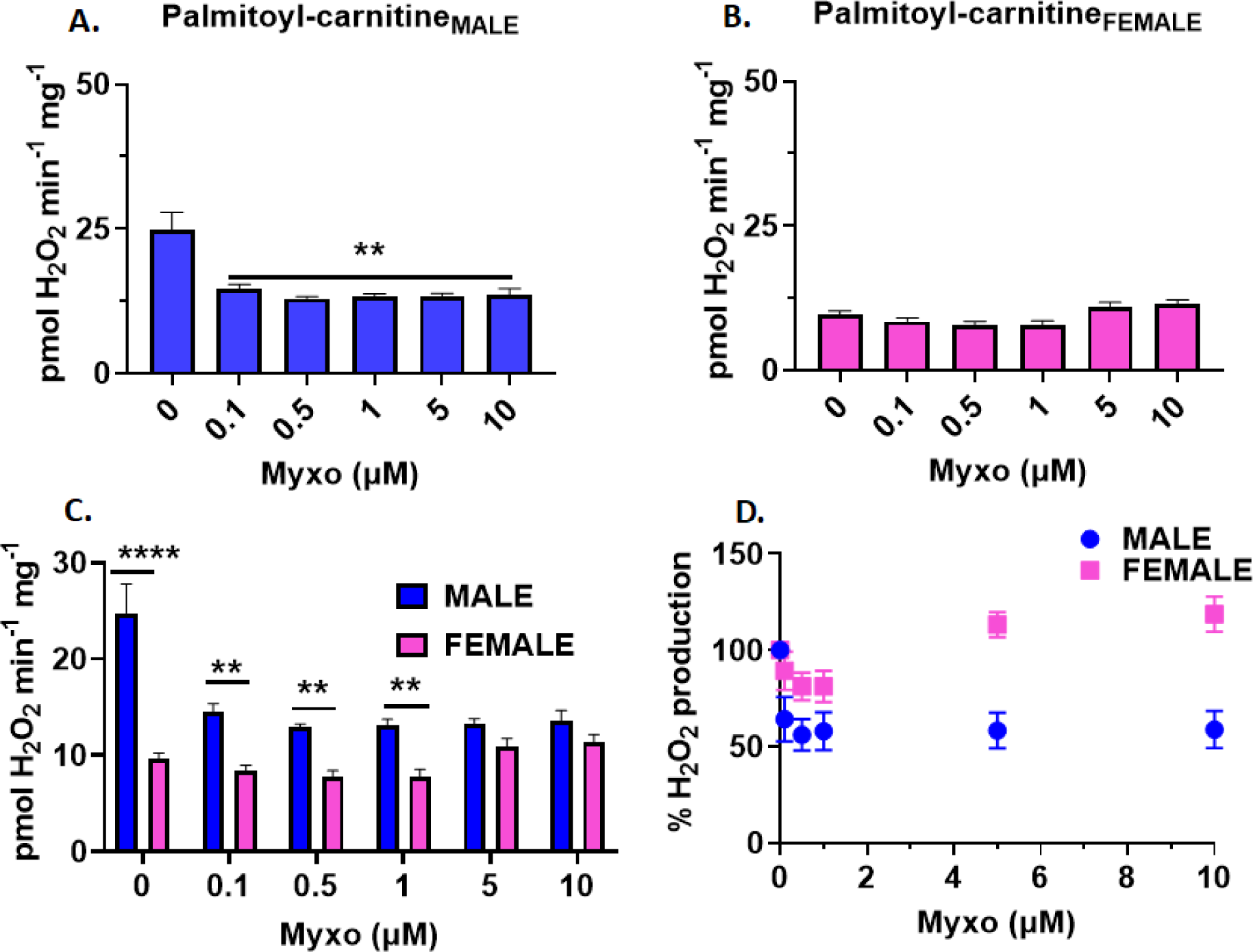
Myxothiazol inhibits palmitoyl-carnitine-induced mH_2_O_2_ production by male, but not female, liver mitochondria. **A.** The effect of increasing doses of competitive and site-specific inhibitor for complex III, myxothiazol, on the rate of mH_2_O_2_ generation by male liver mitochondria oxidizing palmitoyl-carnitine and malate. N=4, mean ± SEM, 1-way ANOV with a post-hoc Fisher’s LSD test. **B.** The effect of increasing doses of competitive and site-specific inhibitor for complex III, myxothiazol, on the rate of mH_2_O_2_ generation by male liver mitochondria oxidizing palmitoyl-carnitine and malate. N=4, mean ± SEM, 1-way ANOV with a post-hoc Fisher’s LSD test. **C.** Comparison of the rates of mH_2_O_2_ production by male and female liver mitochondria energized with palmitoyl-carnitine and malate and incubated in increasing doses of myxothiazol. N=4, mean ± SEM, 1-way ANOVA with a post-hoc Fisher’s LSD test. **D.** Calculation of the percent change in the rate of mH_2_O_2_ production relative to control in response to increasing doses of myxothiazol. N=4, mean ± SEM, 1-way ANOVA with a post-hoc Fisher’s LSD test.

### 2.4. KGDH is a major source of mH_2_O_2_ during palmityol-carnitine oxidation

The results collected above illustrate there are small but significant sex dimorphic effects in the contributions of complexes I, II, and III towards total H_2_O_2_ release from mitochondria. KGDH also exhibited lower rates of H_2_O_2_ generation in female livers when compared to males. Surprisingly, KMV induced ∼90% suppression of H_2_O_2_ generation in both males and females. Notably, inhibition of the ETC only induced small decreases in mH_2_O_2_ generation, suggesting KGDH is the major source of mH_2_O_2_ in liver mitochondria regardless of sex. The results collected above led us to hypothesize KGDH could also be a major supplier of mH_2_O_2_ during fatty acid oxidation. To test this, we supplied mitochondria with palmitoyl-carnitine and malate and then titrated the site specific KGDH inhibito, KMV, into the reaction chambers. **Figure 9A** shows KMV almost abolished mH_2_O_2_ generation when male liver mitochondria were oxidizing palmitoyl-carnitine in the presence of malate. A similar observation was made with the female liver mitochondria **(Figure 9B)**. Indeed, adding KMV to a concentration as low as 0.1 mM induced a significant decrease in mH_2_O_2_ generation **(Figure 9B)**. Notably, in males and females, KMV retained its inhibitory effect at all concentrations provided, even though some variability was detected, especially in the female samples **(Figure 9A and 9B)**. The rate for mH_2_O_2_ generation in mitochondria fueled with palmitoyl-carnitine and malate was lower in the female liver samples **(Figure 9C)**. Calculation of the percent inhibition of mH_2_O_2_ revealed similar trends for the KMV-mediated decrease in production in samples collected from male and female mice **(Figure 9D)**. However, the KMV effect was not as pronounced in the samples collected from the females **(Figure 9D)**.

**Figure 9:**
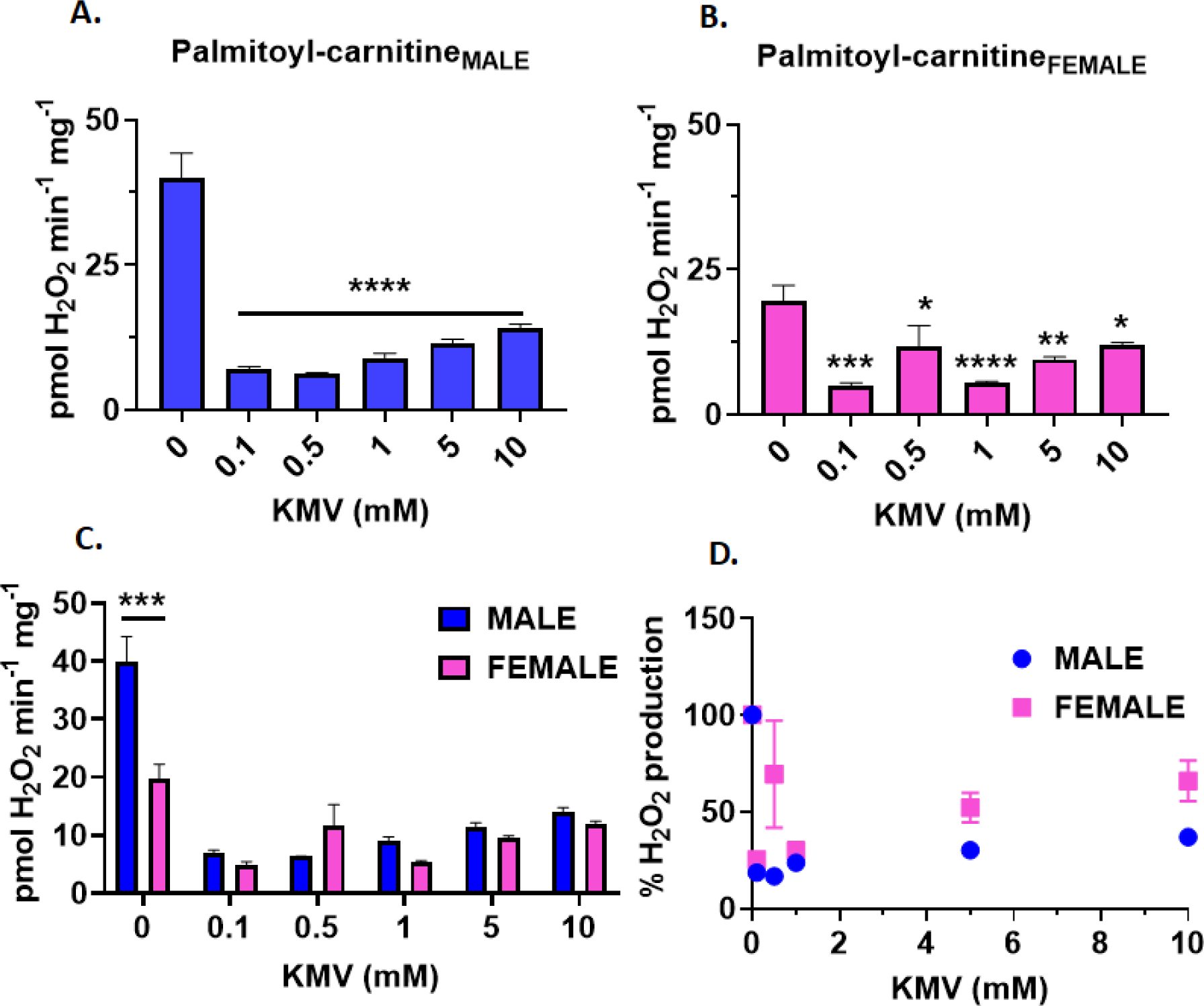
KGDH is a main source of mH_2_O_2_ generation during the oxidation of palmitoyl-carnitine in male and female liver mitochondria. **A.** The effect of increasing doses of competitive and site-specific inhibitor for KGDH, KMV, on the rate of mH_2_O_2_ generation by male liver mitochondria oxidizing palmitoyl-carnitine and malate. N=4, mean ± SEM, 1-way ANOV with a post-hoc Fisher’s LSD test. **B.** The effect of increasing doses of competitive and site-specific inhibitor for KGDH, KMV, on the rate of mH_2_O_2_ generation by female liver mitochondria oxidizing palmitoyl-carnitine and malate. N=4, mean ± SEM, 1-way ANOV with a post-hoc Fisher’s LSD test. **C.** Comparison of the rates of mH_2_O_2_ production by male and female liver mitochondria energized with pyruvate and malate in response and incubated in increasing doses of KMV. N=4, mean ± SEM, 1-way ANOVA with a post-hoc Fisher’s LSD test. **D.** Calculation of the percent change in the rate of mH_2_O_2_ production relative to control in response to increasing doses of KMV. N=4, mean ± SEM, 1-way ANOVA with a post-hoc Fisher’s LSD test.

Next, we examined the impact of S1QEL 1.1 (S1) and S3QEL 2 (S3), inhibitors for mH_2_O_2_ generation by complexes I and III, respectively, on H_2_O_2_ generation in male and female liver mitochondria oxidizing palmitoyl-carnitine in the presence of malate **(Figure 10)**. This was compared to mitochondria treated with KMV, rotenone, or myxothiazol. Note that S1 and S3 interfere with mH_2_O_2_ generation by complex I and III without inhibiting electron conductance by the respiratory chain (Goncalves *et al*, 2020). First, the sex dimorphism in overall mH_2_O_2_ generation was still present with male liver mitochondria exhibiting much higher rates in samples energized with either palmitoyl-carnitine or pyruvate **(Figure 10)**. Surprisingly, S1 and S3, which are known to induce decreased H_2_O_2_ generation by complexes I and III (Goncalves *et al*., 2020), had no effect on the rates of generation in male or female samples oxidizing either palmitoyl-carnitine or pyruvate **(Figure 10)**. By contrast, competitive KGDH inhibitor KMV almost abolished H_2_O_2_ generation regardless of sex or substrate **(Figure 10)**. The addition of KMV to mixtures containing S1 and S3 also almost abolished mH_2_O_2_ production **(Figure 10)**. Rotenone did not induce any significant effects on H_2_O_2_ generation in mitochondria oxidizing pyruvate but did elevate it slightly in samples energized with palmitoyl-carnitine **(Figure 10)**. Myxothiazol suppressed H_2_O_2_ generation in female samples, but not to the same extent as KMV and induced a small but significant decrease in production in the male liver mitochondria. We also examined if palmitoyl-carnitine oxidation alone (e.g., with no malate) could be inhibited by KMV **(Figure S2)**. KMV did inhibit mH_2_O_2_ production, but the effect was not as pronounced as samples metabolizing either pyruvate or palmitoyl-carnitine with malate **(Figure S2)**. We attribute the small KMV effect in the mitochondria treated with palmitoyl-carnitine alone due to presence of endogenous malate in the matrix.

**Figure 10:**
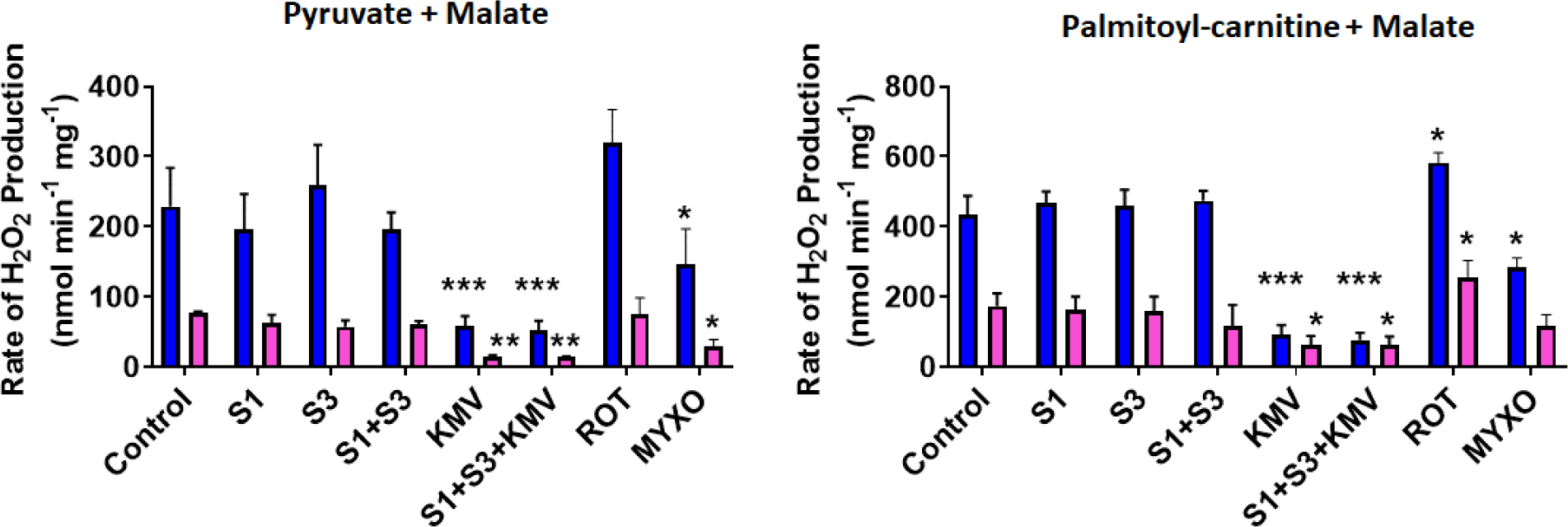
KMV almost abolishes mH_2_O_2_ generation by male and female liver mitochondria whereas complex I and III inhibitors, S1QEL 1.3 and S3QEL 2, have no effect. The rate of mH_2_O_2_ generation was measured in male and female liver mitochondria and after incubation in various complex I and III inhibitors and/or KMV. Complex I inhibitors included S1QEL 1.3 (S1) and rotenone. S1 has been documented to limit mH_2_O_2_ generation while rotenone has the opposite effect. Inhibitors for complex III included S3QEL 2 (S3) and myxothiazol. Both compounds lower mH_2_O_2_ by complex III. N=4, mean ± SEM, 1-way ANOVA with a post-hoc Fisher’s LSD test.

### 2.5 KMV does not inhibit fatty acid oxidation but deactivates mH_2_O_2_ generation induced by medium and short-chained fatty acids

KMV is an α-keto acid that is structurally analogous to α-ketoglutarate. Fatty acid oxidation requires the activity of β-hydroxy acyl-CoA dehydrogenase (HADH), which produces β-keto acyl-CoA for thiolation by CoASH. KMV does have some structural properties that resemble β-hydroxy acyl-CoA. Thus, to eliminate the possibility it was interfering with FAO, we examined its effect on the activity of HADH. Incubation of permeabilized male and female liver mitochondria in KMV did not interfere with the NADH consuming activity of HADH **(Figure 11A)**. KMV also did not disrupt the activity of PDH **(Figure 11B)**. However, KMV did abolish the activity of KGDH in permeabilized liver mitochondria collected from male and female mice **(Figure 11B)**. We also examined if KMV could disrupt succinate-mediated mH_2_O_2_ generation **(Figure 11C)**. KMV inhibited mH_2_O_2_ generation in male and female liver mitochondria supplemented with the pyruvate or palmitoyl-carnitine with malate **(Figure 11C)**. However, it did not interfere with succinate-supported mH_2_O_2_ generation **(Figure 11C)**. Taken together, KMV is specific for KGDH and does not inhibit PDH, FAO, or complex II and the ETC.

**Figure 11:**
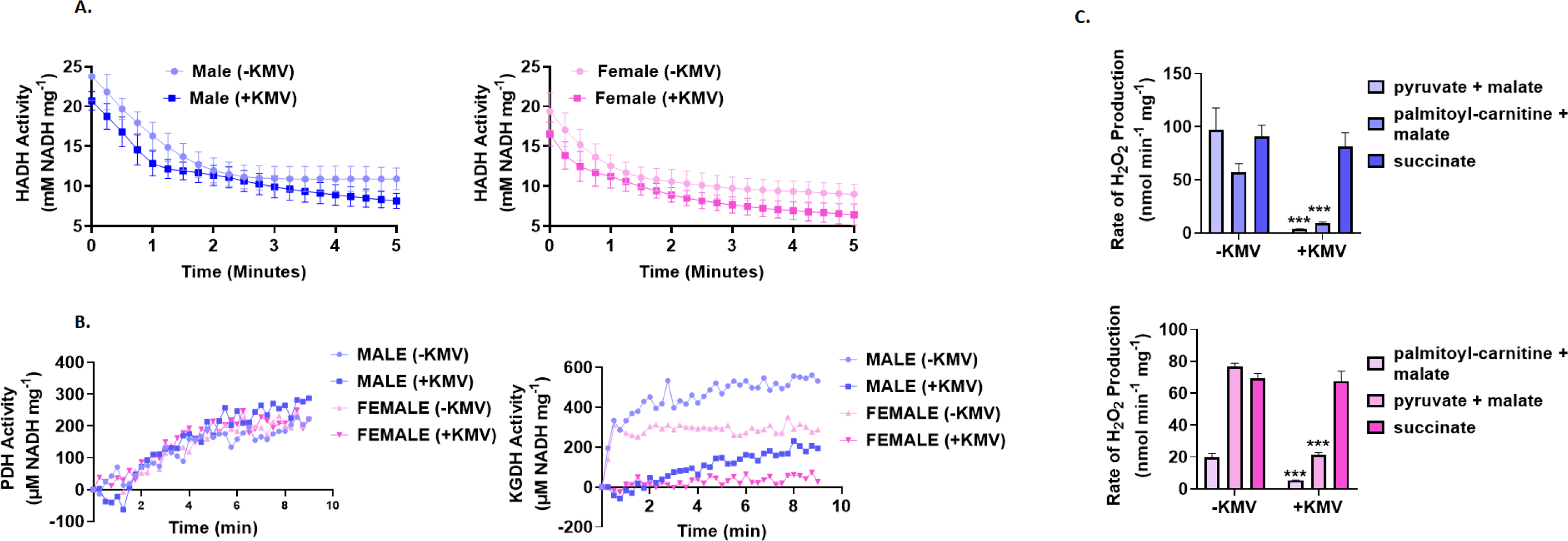
KMV does not interfere with the activity of fatty acid oxidation enzyme, β-hydroxy acyl-CoA dehydrogenase (HADH) or inhibit mH_2_O_2_ generation when succinate serves as the mitochondrial fuel source. **A.** Measurement of the activity of HADH in permeabilized male and female liver mitochondria and the effect of KMV on the reaction. HADH was measured by assessing the consumption of NADH in the presence of acetoacetyl-CoA. N=4, mean ± SEM. **B.** Measurement of the impact of KMV on the activities of pyruvate dehydrogenase (PDH) and α-ketoglutarate dehydrogenase (KGDH) in permeabilized male and female liver mitochondria. N=4, mean ± SEM. **C.** The rate of mH_2_O_2_ generation by male and female liver mitochondria energized with pyruvate and malate, palmitoyl-carnitine and malate, or succinate and treated with our without KMV. N=4, mean ± SEM, 1-way ANOVA with a post-hoc Fisher’s LSD test.

Next, we interrogated the impact of KMV on mH_2_O_2_ generation by male and female liver mitochondria incubated in fatty acyl-carnitines with variable chain lengths. KMV induced a robust inhibition of mH_2_O_2_ generation when palmitoyl-carnitine and malate served as substrates **(Figure 12A and B)**. This inhibition of mH_2_O_2_ generation also occurred with C14 and C8 fatty acyl-carnitines. Indeed, mH_2_O_2_ production was suppressed by the KMV in male and female mitochondria treated with myristoyl-carnitine (C14) and octanoyl-carnitine (C8) and malate **(Figure 12A and B)**. KMV also inhibited mH_2_O_2_ release from male and female liver mitochondria metabolizing butyryl-carnitine (C4) **(Figure 12A and B)**. However, the inhibition was not as robust, especially in the males, which we attribute to butyryl-carnitine being a poor inducer of mH_2_O_2_ generation because of its short chain length.

**Figure 12:**
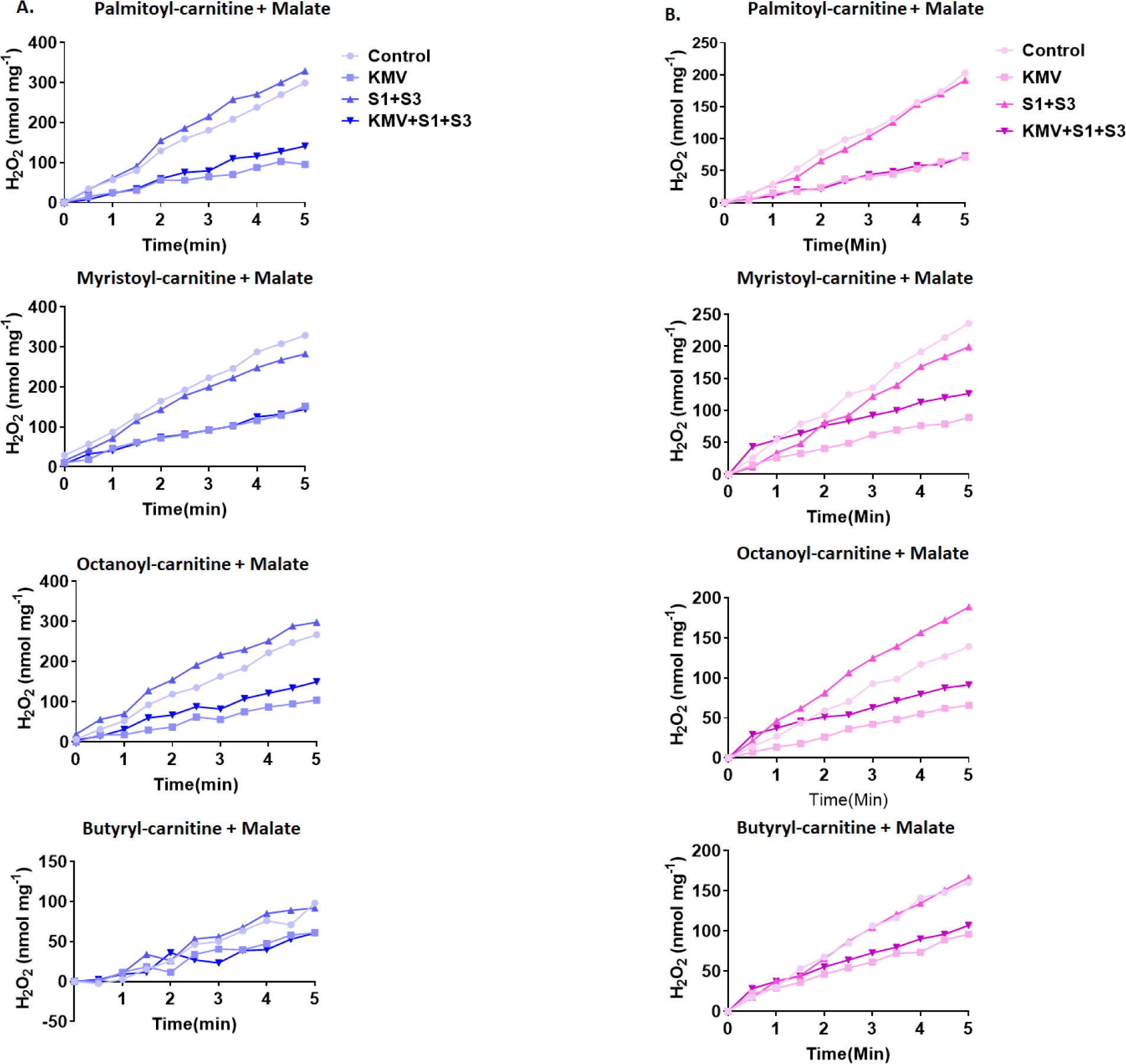
KMV deactivates mH_2_O_2_ generation by male and female liver mitochondria oxidizing long-chain (palmitoyl or myristoyl-carnitine), medium chain (octanoyl-carnitine), short chain (butyryl-carnitine) fatty acids. **A.** Measurement of the rate of mH_2_O_2_ generation by male liver mitochondria oxidizing long, medium, and short-chained fatty acids with malate and in the presence or absence of KMV. N=4, mean ± SEM. **B.** Measurement of the rate of mH_2_O_2_ generation by male liver mitochondria oxidizing long, medium, and short-chained fatty acids with malate and in the presence or absence of KMV. N=4, mean ± SEM.

## 3. Discussion

Complex I and III of the ETC are often viewed as the major, and sometimes only, sources of mH_2_O_2_ in mitochondria. This convention was challenged two decades ago when KGDH was identified as a source of mH_2_O_2_ (Starkov *et al*., 2004; Tretter & Adam-Vizi, 2004). KGDH was originally found to support mH_2_O_2_ production during α-ketoglutarate oxidation in neuronal mitochondria and synaptosomes (Starkov *et al*., 2004; Tretter & Adam-Vizi, 2004). Later efforts revealed KGDH is a main generator in rat skeletal muscle tissue and mouse liver mitochondria (Quinlan *et al*., 2014; Slade *et al*., 2017). To delineate the importance of KGDH in the provision of mH_2_O_2_ in the liver, in the present study we conducted the first ever in-depth assessment of sex dimorphic effects on mH_2_O_2_ production by KGDH and components of the ETC. We discovered sex dimorphisms in mH_2_O_2_ generation by the ETC components, complex I and III, which display differences in production based on sex and bioenergetic conditions. The most notable observation made here is that KGDH was the major source of mH_2_O_2_ production in male and female liver mitochondria, even when fatty acids served as the combustible fuel source. The contribution of fatty acids towards KGDH-driven mH_2_O_2_ generation was dependent on malate priming the Krebs cycle. This is highly novel, considering the ETC is often viewed as the only mH_2_O_2_ source during fatty acid oxidation. Our discovery has strong implications for understanding sex differences in mH_2_O_2_ production and their effect on hepatocyte redox homeodynamics. These findings also implicate KGDH in the activation of oxidative distress and the pathogenesis of MAFLD.

### 3.1 Increased coupling efficiency in OxPhos correlates with the lower rates for mH_2_O_2_ generation by female liver mitochondria

Mitochondria display many sex dimorphisms in bioenergetics and redox poise due to differences in gene expression, protein regulation, and epigenetic coding. Confirmed in this study, female liver mitochondria, in general, produce less mH_2_O_2_ when compared to males (Borras *et al*., 2003). Mitochondria contain 16 sources of mH_2_O_2_, 12 of which are associated with pathways required for the coupling of nutrient oxidation to OxPhos (Goncalves *et al*, 2015). Few studies have investigated the sex effects on these individual sources and their overall contributions towards total mH_2_O_2_ generation. Using different substrate and inhibitor combinations and transgenic mouse models, we had previously shown in several studies PDH and KGDH display rates for mH_2_O_2_ generation that can be up to ∼6-fold higher in male liver mitochondria when compared to the female littermates (Gill *et al*, 2020; Mallay *et al*, 2019; Wang *et al*., 2023). This was also the case for dihydroorotate dehydrogenase (DHODH), an oxidoreductase embedded in the mitochondrial inner membrane that is required for pyrimidine biosynthesis (Koufos & Mailloux, 2023). In the present study, we found rates of mH_2_O_2_ generation were significantly lower in female liver mitochondria when compared to males. Several factors could account for this sex difference in mH_2_O_2_ generation. The concentration of GSH and its corresponding peroxidases (e.g., Gpx1) is greater in female liver mitochondria and hepatocytes (Diaz *et al*, 2019; Valle *et al*., 2007). Additionally, liver mitochondria occur in greater numbers in female rodents, have greater proton leaks, contain a higher mitochondrial DNA copy number and are more enriched in cardiolipin, factors that increase tolerance towards oxidative distress (Fernandez-Vizarra *et al*, 2011; Justo *et al*., 2005). Together, this creates a mitochondrial microenvironment that has greater capacity for mH_2_O_2_ elimination. However, we also found this sex difference was also related to the better coupling efficiency of OxPhos in the female liver mitochondria, which suppressed mH_2_O_2_ generation. Indeed, female liver mitochondria displayed greater ADP and FCCP-stimulated O_2_ consumption, implying a greater number of electrons yielded from pyruvate, succinate, or palmitoyl-carnitine are being successfully delivered to complex IV. This would limit the over-reduction of redox centers in the ETC and Krebs cycle enzymes, lowering the number of electrons available to generate mH_2_O_2_. The increased observed OxPhos efficiency was related to the augmentation of PDH, KGDH, and complex I expression, indicating improved coupling between nutrient oxidation and ATP production is associated with increased expression of the components required for mitochondrial metabolism and electron conductance.

### 3.2 KGDH is the major source of mH_2_O_2_ and thus important for the maintenance of optimal liver health

The surge in MAFLD is related to the increased preponderance of metabolic diseases like obesity, a poor diet, and xenobiotic exposure (Karlsen *et al*, 2022). MAFLD has a wide spectrum of clinical manifestations ranging from simple steatosis to more severe forms like cirrhosis (Gallage *et al*, 2022). A distinguishing feature of MAFLD is intrahepatic lipid accumulation, triggered by abnormal mitochondrial function and defects in fatty acid oxidation (Gallage *et al*., 2022; Ma *et al*., 2023; Zhou *et al*, 2022). This results in mH_2_O_2_ accumulation and the induction of oxidative distress (Kakimoto *et al*, 2021; Kakimoto *et al*, 2015). Unfortunately, the relationship between mH_2_O_2_ availability and liver health and disease is much more complex than simply preventing oxidative distress. Advances in redox biology have identified mH_2_O_2_ as a powerful second messenger that is required for hepatic health through the induction of oxidative eustress pathways. Transient, and controlled bursts in mH_2_O_2_ generation is integral for triggering cell proliferation, survival, and adaptation pathways that promote liver recovery in response to injury (Bai *et al*., 2023; Bai *et al*., 2015). Together, this has generated interest in the therapeutic potential of targeting mitochondria to re-establish homeostatic control over mH_2_O_2_ for the prevention of MAFLD and its progression to mover severe diseases.

Understanding mH_2_O_2_ homeodynamics and its role in maintaining optimal liver health first requires identifying which mH_2_O_2_ sources in mitochondria serve as major suppliers. FAO can generate mH_2_O_2_ by complexes I, II, and III of the ETC (Perevoshchikova *et al*, 2013; Seifert *et al*, 2010). Notably, ETC-independent sources for mH_2_O_2_ production during FAO were suggested by Seifert *et al*, but these generators were never identified (Seifert *et al*., 2010). FAO can drive low rates of mH_2_O_2_ production by electron-transferring flavoprotein oxidoreductase (ETFQO), long-chain acyl-CoA dehydrogenase (LADH), and very long-chain acyl-CoA dehydrogenase (VADH), which donate electrons to the ETC (Kakimoto *et al*., 2015; Perevoshchikova *et al*., 2013; Seifert *et al*., 2010). Thus, the bulk of the mH_2_O_2_ formed during FAO likely does not originate from the ETC or ETFQO, LADH, or VADH. Notably, experiments that estimate mH_2_O_2_ generation by isolated mitochondria fueled with fatty acids are often carried out in the absence of malate since it is assumed the ETC is the main generator. It would be expected that the Krebs cycle is primed during FAO by liver mitochondria *in vivo* since acetyl-CoA units generated by fat degradation need to be oxidized further. We designed our experiments around the concept that the Krebs cycle would be primed during FAO. This led to our discovery KGDH is the major mH_2_O_2_ generator in male and female liver mitochondria mice during FAO, but only when malate was added to reaction mixtures. This rate of production during FAO could almost be abolished by KMV, a site-specific inhibitor for KGDH. KMV elicited a similar inhibitory effect when pyruvate was the energetic substrate, showing PDH is not a main source in liver mitochondria, an observation we had made previously (Oldford *et al*., 2019; Slade *et al*., 2017; Wang *et al*., 2023). We also examined rates of mH_2_O_2_ genesis in the presence of classic complex I and III inhibitors, rotenone and myxothiazol, which increase and decrease, respectively, the rate of production by the ETC **(Figure 1)**. Notably, both compounds only had small but significant effects on the rates of mH_2_O_2_ genesis. In addition, the complex I and III mH_2_O_2_ generation inhibitors, S1 and S3, had little effect on the rates of production. Combining S1 and S3 with KMV, on the other hand, almost abolished mH_2_O_2_ generation when pyruvate or palmitoyl-carnitine were used to energize mitochondria with malate. This finding unequivocally shows KGDH is a main mH_2_O_2_ source. KMV was also effective at suppressing mH_2_O_2_ generation even when medium and short-chained acyl-carnitines served as substrates. Taken in aggregate, our findings demonstrate KGDH is a major mH_2_O_2_ source in liver mitochondria. Furthermore, KGDH occupies a pivotal position in Krebs cycle metabolism since it is the major point of convergence for the further oxidation of acetyl groups and amino acids and is thus ideally positioned to serve as a redox signaling platform for mediating oxidative eustress signals.

### 3.3 KGDH as the major source of hepatic mH_2_O_2_ in the acceleration of oxidative distress in post-menopausal women

Another key observation we made here is there appeared to be no sex dimorphic effect on the importance of KGDH in supplying mH_2_O_2_ in male and female liver mitochondria. This contrasts with complexes I and III which we identified as having sex dimorphisms in their contributions to the total mH_2_O_2_ generated by mitochondria. Our finding that KGDH is the major mH_2_O_2_ source in male and female liver mitochondria has strong implications for understanding the effect of sex on the development of MAFLD and other disorders. MAFLD has a higher preponderance in men when compared to pre-menopausal women (Burra *et al*, 2021). However, MAFLD surges in post-menopausal women, which parallels the onset of obesity, type 2 diabetes mellitus, and other disorders (Carrieri *et al*, 2022). The sex dimorphic effects in MAFLD have been related to superior mitochondrial redox poise, mH_2_O_2_ budgeting, and fuel metabolism in female rodents and pre-menopausal women (Borras *et al*., 2003; Burra *et al*., 2021; Carrieri *et al*., 2022; Justo *et al*., 2005; Nadal-Casellas *et al*., 2010; Valle *et al*., 2007). MAFLD is accelerated in ovariectomized rodents and post-menopausal women due to defects in mitochondria causing oxidative distress (Carrieri *et al*., 2022; Fuller *et al*, 2021). Taken together with our findings, it is likely that KGDH serves as this source of mH_2_O_2_ that triggers oxidative distress and rapidly accelerates the onset of MAFLD in the ovariectomized rodents and in post-menopausal women. Further, the superior redox buffering capacity of livers from female rodents and fertile women would keep the mH_2_O_2_ formed “*in balance*” to promote the activation of beneficial eustress pathways for optimal liver health. Although speculative, targeting KGDH to modulate the rate of its mH_2_O_2_ release may serve as a new way to treat and prevent MAFLD in men, but also post-menopausal women.

Loss of redox poise or control over mH_2_O_2_ generation due to defective mitochondrial metabolism is also an early event in the manifestation of MAFLD and its progression to more severe liver pathologies. Thus, it is vital to decode the details of how mitochondria generate mH_2_O_2_ and the molecular origins of its production since it plays an integral role in maintaining optimal liver health. Here, we have presented evidence showing KGDH is the major mH_2_O_2_ supplier in the liver during the combustion of pyruvate from glycolysis or acetyl-CoA formed from FAO. Overall, this has strong implications for understanding the role of KGDH in triggering hepatic oxidative eustress signals for maintaining liver health and how defects in mH_2_O_2_ production by KGDH causes oxidative distress. More importantly, although there are important sex dimorphic effects on overall mH_2_O_2_ biogenesis, we showed for the first time KGDH is the major source for this mH_2_O_2_ regardless of sex. These findings point to the importance of this site in the sex-independent activation of oxidative eustress signals and illustrates that KGDH could be targeted for the treatment of MAFLD in men and post-menopausal women.

## 4. Material and Methods

### 4.1 Animals and preparation of mouse liver mitochondria

Animals were cared for in accordance with the principles and guidelines of the Canadian Council on Animal Care (CCAC) and the Institute of Laboratory Animal Resources. Animal experiments were approved by the Facilities Animal Care Committee (FACC) in the Faculty of Agriculture and Environmental Sciences at McGill University. Wild-type male and female C57BL/6N mice aged 9 weeks were purchased from Charles River Laboratories. They were housed at 25℃ on a 12-hour day/night light cycle and provided unlimited access to water and standard chow (TD2020X; Inotiv).

Mitochondria isolated from mice ranging from 10 to 13 weeks were utilized in experiments. Mice were euthanized via cervical dislocation while under 5% isoflurane. Livers were surgically removed and placed in cold MESH buffer (220 mM mannitol, 1 mM EGTA, 70 mM sucrose, 10 mM HEPES, pH 7.4). All steps were performed on ice and/or at 4 °C. Mouse livers were cut into pieces and washed in ice-cold MESH buffer to remove excess blood and hair. The livers were minced using a razor on a Teflon watch glass and then homogenized in 25 mL of MESH supplemented with 0.5% w/v defatted BSA using the Potter-Elvejham method and variable speed homogenizer (Glas-Col). Homogenates were centrifuged at 900 x g for 9 min to pellet cellular debris and nuclei (Sorvall Lynx 4000). The supernatant was collected and centrifuged at 12000 x g for 9 mins to pellet the mitochondria. The pellet was resuspended in 500 μL of MESH and the protein equivalents were determined using a Bradford assay utilizing a BSA standard.

### 4.2 mH_2_O_2_ production assays

The rate of mH_2_O_2_ production by isolated mitochondria was measured using the Amplex UltraRed (AUR) assay and our established substrate/inhibitor toolkit (reviewed in (Grayson & Mailloux, 2023)). Isolated mitochondria were diluted to 5 mg/mL in MESH and then stored on ice prior to assays. Samples were then diluted to 0.5 mg/ml in MESH in the wells of a 96-well black plate. The final reaction volume was 200 µL. The mitochondria were allowed to equilibrate at 25 °C for ∼5 min and were then treated with the following inhibitor and substrate combinations: 1) succinate (5 mM) with malonate (0-100 mM; complex II inhibitor), 2) 5 mM pyruvate and 2 mM malate with KMV (0-10 mM), 3) 100 µM palmitoyl-carnitine and 5 mM malate with rotenone (1 µM) and myxothiazol (0-10 μM), and 4) 100 μM palmitoyl-carnitine and 2 mM malate with rotenone (1 µM), myxothiazol (5 µM) and KMV (0-10 mM). In certain cases, only palmitoyl-carnitine, myristoyl-carnitine, octanoyl-carnitine, or butyryl-carnitine or pyruvate with malate or succinate was added with KMV, rotenone, or myxothiazol. A full breakdown for the sites of action for the different substrate and inhibitor combinations is supplied in **Figure 1**. The mitochondria were then supplemented with horseradish peroxidase (HRP; 30 U/mL), superoxide dismutase (SOD; 25 U/mL), and Amplex UltraRed (AUR; 20 uM). The conversion of Amplex UltraRed to fluorescent resorufin was measured to determine mH_2_O_2_ production. The kinetic measurement was recorded every 30 sec for 5 minutes at 565/600 nm using a Cytation 5 microplate reader equipped with Gen 5 3.11 software. The production of mH_2_O_2_ was normalized to mitochondrial protein equivalents and background fluorescence. The rate of mH_2_O_2_ production was calculated using a linear standard curve for the reaction of 1-1000 mM mH_2_O_2_ with AUR and HRP.

### 4.3 Seahorse XFe24 Assays

Examination of mitochondrial bioenergetics using the XFe24 was performed as described previously (Mailloux *et al*, 2013). Briefly, mitochondria were isolated from livers as described above except the MESH contained 10 mM pyruvate and 2 mM malate. Samples were diluted to 0.2 mg/mL and 50 µL was added to the wells of a XFe24 tissue culture plate. The plate was then centrifuged at 1200 xg for 20 min at room temperature in a swing bucket centrifuge (Sorvall X Pro Series). The wells were then supplemented with 450 µL of respiration buffer (MESH supplemented with 10 mM KH_2_PO_4_, 2 mM MgCl_2_, and 0.2% (w/v) defatted BSA and pyruvate (10 mM)/malate (2 mM), succinate (5 mM), or palmitoyl-carnitine (100 µM)/malate (2 mM)). The mitochondria were then incubated in the buffer for 30 min at 37 °C. Injection protocols were developed on an Agilent Seahorse XFe24 Analyzer using Wave Controller Software 2.4. The mitochondrial oxygen consumption rate (OCR) under the different states of respiration was interrogated by first measuring state 4 respiration (substrates alone with mitochondria). State 3, state 4_O_ (oligomycin), and state 4_U_ (uncoupled) were induced by injecting ADP (1 mM), oligomycin (2.5 μg/mL), and FCCP (4 μM), respectively. Antimycin A (4 μM) was injected at the end of the XFe24 protocol to arrest the electron transport chain. OCR values were normalized to mitochondrial content and rates of O_2_ consumption after the antimycin A injection.

### 4.4. β-hydroxyacyl-CoA dehydrogenase (HADH) assay

HADH was assayed using permeabilized mitochondria. Briefly, mitochondria were diluted to 1 mg/mL in MESH containing 0.1% triton x-100 and then incubated on ice for 60 min. Samples were then diluted to 0.1 mg/mL in MESH, equilibrated for 5 min, and then incubated for 15 min in KMV (10 mM). Acetoacetyl-CoA (0.09 mM) and NADH (0.1 mM) were then added. The final reaction volume was 200 µL. HADH activity was estimated by measuring NADH consumption at 340 nm. The activity of the enzyme was calculated using a molar extinction coefficient of ε_340_ = 6220 M^-1^cm^-1^ and the Beer-Lambert Law.

### 4.5 PDH and KGDH activity assays

PDH and KGDH were assayed essentially the same way as HADH except the activities were estimated by measuring NADH production. Permeabilized mitochondria diluted to 0.1 mg/mL in MESH and incubated in KMV (10 mM). CoASH (0.1 mM), TPP (0.3 mM), NAD (0.1 mM), and either pyruvate or α-ketoglutarate (0.1 mM) were then added to each.

### 4.6 Immunoblot

Mitochondria were diluted to 2 mg/mL in 1 x Laemmli buffer containing RIPA solution and then incubated for 10 min at 100 °C. 20 µL of sample was loaded in each well and proteins were electrophoresed through a 10% isocratic denaturing acrylamide gel. Proteins were then electroblotted to nitrocellulose membranes by tank transfer. Successful transfer was confirmed by Ponceau S staining. Membranes were blocked for 1 hour with TBS-T containing 5% (w/v) non-fat skim milk (blocking solution), washed twice with TBS-T, and then incubated overnight in primary antibodies directed against PDH (Abcam, PDH cocktail), KGDH (Abcam, E1 subunit), and the respiratory chain (Abcam, OxPhos cocktail). These primary antibodies were diluted in TBS-T containing 5% (w/v) defatted BSA and 0.005% NaN_3_. The dilution factor for all primary antibodies used was 1/1000. Membranes were then washed thrice with TBS-T and probed with secondary goat anti-mouse and anti-rabbit tagged with HRP (Abcam). The secondary antibodies were diluted in blocking solution and it was a 1/2000 dilution factor. Activity bands were visualized with chemiluminescent substrate (Thermofisher) and imaged using a Licor C-Digit Scanner (Licor). Bands were quantified using ImageJ software.

### 4.7 Statistical Analysis

Rates of mH_2_O_2_ generation by liver mitochondria, enzyme activities, and XFe24 data were processed in EXCEL. Rate results and traces were then collated into Graphpad Prism 9 for statistical analyses. XFe24 assays were performed in quadruplicate from three different samples. OCR results from XFe24 assays were collected using Wave Controller Software 2.4 (Agilent). Analysis of immunoblot data was conducted on three separate samples. mH_2_O_2_ generation assays and enzyme activities were conducted in duplicate and on at least four different samples. Rates for mH_2_O_2_ generation were estimated using standard curves. Results were analyzed by unpaired Student T-test or one-way or two-way ANOVA with a Fisher’s least significant square post-hoc test. * = p≤0.05, ** = p≤0.01, *** = p≤0.005, **** = p≤0.001. Fig 1 was generated using Biorender software.

## Acknowledgements

Funding was obtained from the Natural Sciences and Engineering Research Council (NSERC) of Canada Discovery Grant Program (RGPIN-2022-03240) and Start-Up funds provided by McGill University. CG received a Canada Graduate Scholarship – Masters (CGS-M) via the Canadian Institutes of Health Research (CIHR). BF and OK were funded by the Natural Sciences and Engineering Research Council (NSERC) USRA program.

Conception of Project: RJM. Funding: RJM. Figures: Fig 1 (OK, CG), Fig 2 (CG, RJM), Fig 3 (CG, BF, RJM), Fig 5 (OK, CG), Fig 6 (OK, CG), Fig 7 (OK, CG), Fig 8 (OK, CG), Fig 9 (OK, CG), Fig 10 (BF, CG), Fig 11 (BF, CG), Fig 12 (BF, CG), supplemental figures (CG, BF). Experimental design: CG and RJM. Manuscript writing and editing: CG and RJM

